# Characterising the contribution of auditory and somatosensory inputs to TMS-evoked potentials following stimulation of prefrontal, premotor and parietal cortex

**DOI:** 10.1101/2023.11.09.566371

**Authors:** Mana Biabani, Alex Fornito, Mitchell Goldsworthy, Sarah Thompson, Lynton Graetz, John G. Semmler, George M. Opie, Mark A. Bellgrove, Nigel C. Rogasch

## Abstract

Transcranial magnetic stimulation (TMS) results in a series of evoked potentials (TEPs) in electroencephalography (EEG) recordings. However, it remains unclear whether these responses reflect neural activity resulting from transcranial stimulation of the cortex, or from the sensory experiences of TMS. Across three experiments (total n = 135), we recorded EEG activity following TMS to the dorsolateral prefrontal cortex, premotor cortex, and parietal cortex as well as a sensory control condition (stimulation of the shoulder or electrical stimulation of the scalp with a click sound). We found that TEPs showed a stereotypical frontocentral N100/P200 complex following TMS of all cortical sites and control conditions, regardless of TMS intensity or the type of sensory control. In contrast, earlier TEPs (<60 ms) showed site-specific characteristics which were largest at the site of stimulation. Self-reported sensory experiences differed across sites, with prefrontal stimulation resulting in stronger auditory (click sound perception) and somatosensory input (scalp muscle twitch, discomfort) than premotor or parietal stimulation, a pattern that was reflected in the amplitude of later (N100/P200), but not earlier (<60 ms) TEP peak amplitudes. Later TEPs were also larger in individuals who experienced stronger click sound perception and, to a lesser extent, TMS-evoked scalp muscle twitches. Increasing click sound perception by removing auditory masking increased N100/P200 amplitudes without altering earlier peaks, an effect which was more prominent at sites with more successful masking. Together, these findings suggest that the frontocentral N100/P200 complex represents a generalised sensory response resulting from TMS-related auditory and somatosensory input. In contrast, early TEP peaks likely reflect activity resulting from transcranial stimulation of the cortex. The results have important implications for designing and interpreting TEP studies, especially when comparing TEPs between stimulation sites and participant groups showing differences in sensory experiences following TMS.

## 1. Introduction

Transcranial magnetic stimulation (TMS) is a non-invasive brain stimulation technique that has become increasingly utilised in both experimental and clinical neuroscience. A single TMS pulse induces a set of time-locked deflections in electroencephalographic (EEG) recordings of cortical activity, known as TMS-evoked EEG potentials (TEPs) (Ilmoniemi and Kičić, 2010; Rogasch and Fitzgerald, 2013). TEPs reflect neuronal reactivity to TMS, both at the site of stimulation and across the brain, and are sensitive to changes in stimulation parameters (Casarotto et al., 2010), pharmacological manipulation (Premoli et al., 2014), and differences in brain state (Massimini et al., 2005). Furthermore, certain characteristics of TEPs differ across the lifespan and with various neurological and psychiatric disorders (e.g., epilepsy and schizophrenia) (Hui et al., 2019; Tremblay et al., 2019). Despite their promise for understanding human neurophysiology, TEPs are yet to be used as diagnostic markers in clinical contexts as their underlying neurophysiological mechanisms are not well-understood (Tremblay et al., 2019). One crucial criterion for TEPs to be considered as indices of cortical excitability is their sensitivity to the stimulation of different neuronal subsets. However, recent studies have shown that stimulation of different cortical areas, with different neuronal compositions, produce TEP components strongly resembling those induced by sensory sham stimulation (which only produced TMS scalp sensations and/or click sound) (Biabani et al., 2019; Conde et al., 2019; Gordon et al., 2018). These findings have raised serious concerns about whether TEPs reflect dynamical properties of the stimulated cortical circuits or just contain stereotypical sensory responses.

Given these concerns, a growing body of research has examined the contribution of multisensory potentials to TEPs (Biabani et al., 2021, 2019; Conde et al., 2019; Freedberg et al., 2020; Gordon et al., 2023, 2021, 2018; Rocchi et al., 2021). Studies comparing TEPs between stimulation sites have shown that while early latency responses within the first 60 ms following TMS tend to show site-specific topographies and time courses, later responses often converge on a common frontocentral N100 and P200 peak regardless of stimulation site (Freedberg et al., 2020; Rogasch et al., 2020). The spatial and temporal characteristics of these responses are consistent with auditory evoked potentials (AEPs) resulting from the TMS clicking sound (Nikouline et al., 1999), and somatosensory evoked potentials (SEPs) resulting from stimulation of cranial/facial nerves and scalp muscles across the scalp (Gordon et al., 2021), which can cause sensations of discomfort or even pain. The multisensory nature of the frontocentral N100/P200 components has led to the term peripherally-evoked potentials (PEPs). While the contribution of AEPs to TEPs has been recognised for several decades (Nikouline et al., 1999), it was generally assumed that the influence of these potentials could be sufficiently minimised by playing continuous masking noise through headphones and placing a layer of foam between the scalp (Ilmoniemi et al., 2015). However, recent research on TEPs from motor and non-motor cortex has shown these commonly used masking methods are not always sufficient to prevent perception of the coil click, with strong correlations between TEPs and PEPs still often present between 60-300 ms after the TMS pulse (Biabani et al., 2019; Gordon et al., 2023), and in one study as early as 25 ms (Conde et al., 2019). Furthermore, there is ongoing debate as to the relative contribution of AEPs and SEPs to PEPs, with some studies arguing that PEPs are primarily auditory (Ilmoniemi and Kičić, 2010; Paus et al., 2001; Rocchi et al., 2021), while others have suggested a combination of both inputs (Conde et al., 2019; Gordon et al., 2021). Given that somatosensory experiences such as discomfort and scalp muscle twitches differ across scalp locations (Meteyard and Holmes, 2018), it is possible that the relative auditory/somatosensory contribution to TEPs could also differ between stimulation locations. Consequently, methods developed for suppressing sensory input in one region may not always translate to others. Understanding the contribution of sensory inputs to TEPs from different stimulation locations is therefore paramount for interpreting the neural basis of TMS-evoked EEG activity, and for designing effective methods to improve the reliability of TMS-EEG research across the field.

The aim of the current study was to characterise the contribution of auditory and somatosensory inputs to TEPs following stimulation of three non-motor brain regions, the prefrontal cortex, premotor cortex and parietal cortex. In experiment A, we compared TEPs following stimulation of each site to PEPs from TMS over the shoulder, a non-cortical sensory control condition. We also compared self-reported sensory experiences from each site including auditory (loudness of the TMS click) and somatosensory (discomfort, pain, scalp muscle twitch) inputs. In experiment B, we replicated the findings of experiment A in a larger independent cohort and assessed the robustness of these findings to stimulation parameters by using a lower relative stimulation intensity. We then compared individuals with stronger/weaker auditory and somatosensory experiences following stimulation to assess whether the different sensory domains independently altered TEP characteristics. To address limitations with the correlational and between-subject analyses in experiments A and B, in experiment C we performed within-subject manipulations of the level of auditory masking for each stimulation site to more directly assess the role of auditory input for TEPs at each site. We also compared TEPs from each stimulation site with PEPs from a more realistic somatosensory control condition (electrical scalp stimulation matched for each site including a concomitant coil click). We hypothesised that the amplitude of common frontocentral N100 and P200 peaks would be modified by varying levels of both auditory and somatosensory input within and between subjects for each site and between sites, whereas earlier site-specific peaks would remain independent of sensory input.

## 2. Methods

### 2.1 Participants

Experiments A and B were conducted at Monash Biomedical Imaging, Monash University and recruited 29 (17 female, Mean ± SD_Age_: 34 ± 7.25 years) and 94 (58 female, Mean ± SD_Age_ : 27.04 ± 7.01 years) healthy adults; respectively. Experiment C recruited 12 healthy individuals (eight female, Mean±SD_Age_ : 23 ± 4.30 years) and was conducted at the Neurophysiology of Human Movement Laboratory, University of Adelaide. The studies were approved by Monash University (Experiment A-B) and University of Adelaide (Experiment C) Human Ethics Committees and all participants provided their written informed consent prior to participation. Experiments were conducted in accordance with the Declaration of Helsinki and all procedures within the TMS safety guideline of the International Federation of Clinical Neurophysiology were followed (Rossi et al., 2009). During TMS, participants were seated comfortably in an armchair with their eyes open looking straight ahead at a black screen.

### 2.2 EMG

Electromyographic (EMG) activity was measured from the right first dorsal interosseous (FDI) muscle to estimate resting motor threshold. A pair of bipolar surface electrodes (Ag-AgCl with 4mm active diameter) were placed in a belly-tendon montage with a distance of ∼2 cm. The ground electrode was positioned over the midpoint of the middle metacarpal bone in the right hand. EMG signals were sampled at 5kHz, amplified 1000 times, band-pass filtered between 10 and 1000 Hz and epoched between -200 to 500 ms around the TMS pulse.

### 2.3 EEG

EEG was recorded with a TMS-compatible 62-channel SynAmps^2^ EEG system (Neuroscan, Compumedics, Australia) in Experiments A and B and an eego^tm^ mylab device (ANT Neuro, Enschede, The Netherlands) in Experiment C. The equipment used Ag/AgCl-sintered ring electrodes positioned in an elastic cap according to the 10-10 international system (EASYCAP, Germany). The recordings were online-referenced to FPz and grounded to AFz. For each individual, the 3-D positions of all electrodes were digitised and co-registered to their T1-weighted MRI scan using a neuronavigation system (Brainsight™ 2, Rogue Research Inc., Canada). EEG signals were sampled at 10kHz, amplified 1000 times, band-pass filtered between DC and 2KHz, and recorded by the Curry8 (Neuroscan, Compumedics, Australia; Experiments A, B) and eego (Version 1.9.2, ANT Neuro Corp, The Netherlands; Experiment C) software. Impedance at all channels was kept at below 5 kΩ throughout the session.

### 2.4 TMS

Experiments A and B involved a MagPro X100 Option stimulator (MagVenture, Denmark) with a figure-of-eight coil (C–B60), which was set to deliver single TMS pulses with a biphasic waveform. The direction of the electrical current induced in the underlying cortex was anterior-posterior and then posterior-anterior relative to the coil handle. Experiment C used a Magstim 200^2^ stimulator with a figure-of-eight coil (external diameter of each wing 90 mm) delivering monophasic pulses in posterior-anterior direction. The stimulators were controlled by the MATLAB-based MAGIC (MAGnetic stimulator Interface Controller) Toolbox (Saatlou et al., 2018).

Motor hotspot was determined as the position and orientation of the coil that elicited largest motor evoked potentials (MEPs) at a slightly suprathreshold stimulation intensity. Resting motor threshold (rMT) was defined as the minimum intensity of stimulation over the Motor hotspot, with EEG cap on, producing MEPs >50 μV in FDI muscle in at least five out of ten successive trials. Each individual received four blocks of 120 trials at the intensities of 120% (Experiment A, C) or 100% rMT (Experiment B); accounting for coil-to-cortex distance (Julkunen et al., 2009; Stokes et al., 2005). In each block, TMS pulses were delivered with 4-6 seconds intervals (jittered) over one of the targeted cortical sites defined by MNI coordinates (x, y, z), including prefrontal (MNI coordinates: -46, 33, 34), premotor (MNI coordinates: -23, 4, 63) and parietal (MNI coordinates: -26, -69, 63). In addition, shoulder stimulation over the left acromioclavicular joint was added as a control condition to produce PEPs without transcranially activating the cortex. Although shoulder stimulation cannot serve as an optimal control condition, our previous study clearly showed that it produces a substantial amount of PEP signals in TEPs (Biabani et al., 2019). The intensity for shoulder stimulation was adjusted subjectively to the perceived intensity of the strongest stimulation required over the scalp.

To account for inter-individual variability in gyrification of the cortex (Thielscher et al., 2011), the anatomical locations of the cortical sites were verified on the T1-weighted MRI for each individual and coordinates found deep in the sulcus were moved to the adjacent gyrus. The coil was positioned perpendicular to the long axis of the target gyrus to induce the strongest e-field within the area (Thielscher et al., 2011). The MR-based neuronavigation system ensured precise placement and maintenance of TMS coil across trials. To minimise PEPs, a thin layer of foam was attached under the coil and white noise was played through inserted headphones fitted inside disposable foam earplugs worn by participants. To adjust the white noise pressure for each individual, we gradually increased the sound level until the participant was unable to hear the click sound of the TMS coil when discharged at the maximum intensity required across the cortical stimulation conditions or when their upper limit of comfort was reached. The order of the stimulation conditions was counterbalanced for each participant. After each block of stimulation, the participant was asked to score their perception of stimulation on a visual analogue scale (VAS) ranging from 0 to 10 in terms of : i) discomfort (0 = not uncomfortable at all; 10 = highly uncomfortable), ii) pain (0 = no pain at all; 10 = the worst pain I could tolerate during the experiment), iii) muscle twitch (0 = no twitches; 10 = very strong cramp), and iv) click sound (0 = I could not hear the pulses at all; 10 = the pulses were as loud as without white noise).

Within experiment C, an auditory control condition (TMS without noise masking) and a realistic sham condition were also applied to each of the three cortical sites, resulting in six additional blocks of stimulation. For the sham condition, an electrical stimulation (ES) was delivered using a DS7A electric stimulator (Digitimer Ltd., Ft. Lauderdale, Florida, USA) through bipolar electrodes (diameter: 8mm; height: of 3.6 cm) positioned at the site of stimulation over the EEG cap with a distance of 30 mm between the electrodes. Stimulation was delivered as square pulses with the duration of 1000 μs and a maximum compliance voltage of 400V, with the timing synchronised to TMS pulses. According to previous studies, these settings produce an ES below the level required to directly activate the cortex (Cohen and Hallett, 1988; Merton and Morton, 1980). For each stimulation site, the intensity of the ES was adjusted to match the sensation induced by real TMS reported by each individual. All other aspects of the ES block were consistent with the real TMS condition except that the TMS coil was tilted at 90° with respect to the scalp to produce the clicks without directly stimulating the cortex.

### 2.5 Data Analysis

For experiment A, EEG data were pre-processed adopting the method described in (Rogasch et al., 2017, 2014) with custom scripts written in MATLAB (R2020b, The Mathworks, USA) using functions implemented in EEGLAB (Delorme and Makeig, 2004) and TESA (Rogasch et al., 2017) toolboxes. Cortical sources of the evoked potentials were estimated using the Brainstorm (v3) software (Tadel et al., 2011) and customised MATLAB scripts using minimum norm estimation (MNE) method. The scalp EEG cleaning pipeline and the MNE method used in the present study have been described in detail in our recent publication (Biabani et al., 2019) and the scripts can be downloaded from (https://github.com/BMHLab/TEPs-PEPs). The distribution of TMS-induced e-field in the brain was estimated using the SimNIBS software pipeline (www.simnibs.org). First, the individualised head models were reconstructed from T1- and T2-weighted MRIs utilising FreeSurfer (Fischl, 2012) and FSL (Jenkinson et al., 2012) tools. The models consisted of a tetrahedral volume mesh divided into five different tissue types (skin, bone, cerebrospinal fluid, white and grey matter) with default isotropic conductivity values (Thielscher et al., 2011; Windhoff et al., 2013). Following segmentation and meshing, the finite element method (FEM) was used to estimate e-field values. For each cortical region, individualised e-field maps were generated using the stimulation intensity, time-varying current in the coil (dI/dt(A/s)), MNI coordinates of the coil centre and the orientation of the coil in space. Both source and e-field maps of each individual were transformed to the default FSAverage surface template parcellated with Desikan-Killiany atlas (Desikan et al., 2006) for group analysis. Global mean field power (GMFP) was calculated using the method introduced by (Lehmann and Skrandies, 1980),

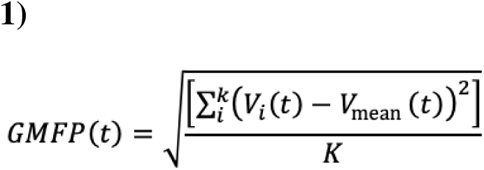

where t is time, K is the number of channels, Vi is the voltage at channel i, and Vmean is the mean voltage across all channels.

For experiments B and C scalp EEG data were pre-processed using the same method as adopted in experiment A. However, the ES condition in Experiment C had one additional pre-processing step for removing the decay artefact using the method employed by (Conde et al., 2019). In this method the exponential model *A*exp(B*x) + C*exp(D*x)* was fitted to the raw data where x was the time between 5 and 500 ms post-stimulus. The parameters A, B, C and D were estimated using fit() function in MATLAB with exp2 as argument. The best fit was then subtracted from each trial and each channel to remove the decay.

All statistical tests were performed in MATLAB using custom scripts. Normality of data distribution was tested using Shapiro-Wilk normality tests. Comparisons across conditions were conducted using One-way ANOVA or Kruskal-Wallis tests depending on the distribution, followed by false discovery rate (FDR)-corrected pairwise comparisons. Spatiotemporal correlations between the potentials among the stimulation conditions were examined using Spearman’s correlation coefficients. The correlation tests were first performed at the individual level and then the coefficients (ρ) were transformed to z values employing Fisher’s transformation for group level statistics (Dunn and Clark, 1969; Hopkins, 1998). The z-scores were tested against zero using one-sample permutation tests with 10000 iterations and t_max_ correction to evaluate the statistical significance of correlations (Blair and Karniski, 1993). For illustration purposes we transformed z back to ρ values.

## 3. Results

### 3.1 Experiment A

All the measurements were well-tolerated by all participants without any serious adverse effects. Figure 1 provides the individual values of the anatomical measures and stimulation parameters for experiment A. Coil-to-cortex distance was significantly different across stimulation sites (one-way ANOVA : F (2, 84) = 7.03, p = 0.001) with the smallest distance at prefrontal cortex and largest distance at premotor cortex (two-sample t-tests FDR-corrected; prefrontal-premotor, p = 0.002; prefrontal-parietal, p = 0.005; premotor-parietal, p = 0.36). Accordingly, prefrontal cortex received the weakest and premotor cortex received the strongest stimulation intensity (one-way ANOVA : F(2, 84) = 4.70, p = 0.011; two-sample t-tests FDR-corrected; prefrontal-premotor, p = 0.007; prefrontal-parietal, p = 0.14; premotor-parietal, p = 0.14). Despite adjusting the stimulation intensity for coil-to-cortex distance, the strength of the generated E-field at the level of cortex was different across stimulation sites (one-way ANOVA : F(2, 84) = 4.86, p = 0.010; two-sample t-tests FDR-corrected; prefrontal-premotor, p = 0.09; prefrontal-parietal, p = 0.27; premotor-parietal, p = 0.008) suggesting the commonly used coil-to-cortex correction method is insufficient for matching TMS-evoked E-fields between sites (Figure 1).

**Figure 1.**
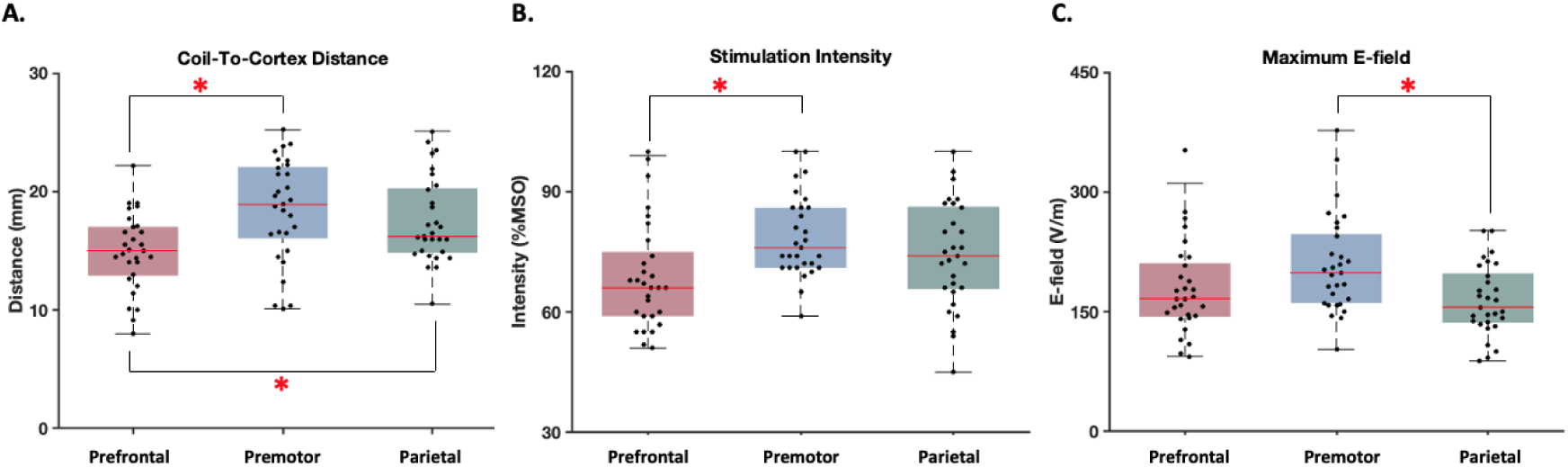
Anatomical measures and stimulation parameters. Each dot in the box and whisker plots represents the value for each individual. The shaded boxes highlight the 25th to 75th centiles of the values and the red horizontal lines within the boxes show the median of the values. **\***indicates significant difference between the stimulation conditions (FDR-corrected p<0.05).

#### 3.1.1 Self-reported sensory experiences of TMS pulses

Figure 2 shows comparisons between sensory experiences following TMS pulses across the different stimulated regions in experiment A. According to Shapiro-Wilk normality tests, VAS values were not normally distributed in all conditions (p>0.05). Therefore, we used Kruskal-Wallis tests to compare each sensation across the stimulation conditions, followed by FDR corrected Wilcoxon Signed-Rank tests for pairwise comparisons. Comparing somatic sensations, parietal cortex stimulation resulted in the lowest ratings of muscle twitch compared to all the other sites (Chi-square = 18.19, df = 3, p = 0.0004; parietal-prefrontal, p = 0.03; parietal-premotor, p = 0.04; parietal-control, p = 0.04) and tended to show the lowest ratings of pain (Chi-square = 6.84, df = 3, p = 0.077) and discomfort (Chi-square = 6.66, df = 3, p = 0.083) across cortical sites, although neither of these reached the threshold for statistical significance. Of note, nearly all conditions reported some perception of the TMS click sound, even though white-noise was played through in-ear headphones in an attempt to mask the click sound. Despite having the lowest stimulation intensity, TMS over prefrontal cortex resulted in the strongest perception of the click sound compared to other sites (Chi-square for sound = 8.35, df = 3, p = 0.04; prefrontal-premotor, p = 0.015; prefrontal-parietal, p = 0.015; prefrontal-control, p = 0.18). These findings show that the sensory experiences differ between stimulation sites across the scalp.

**Figure 2.**
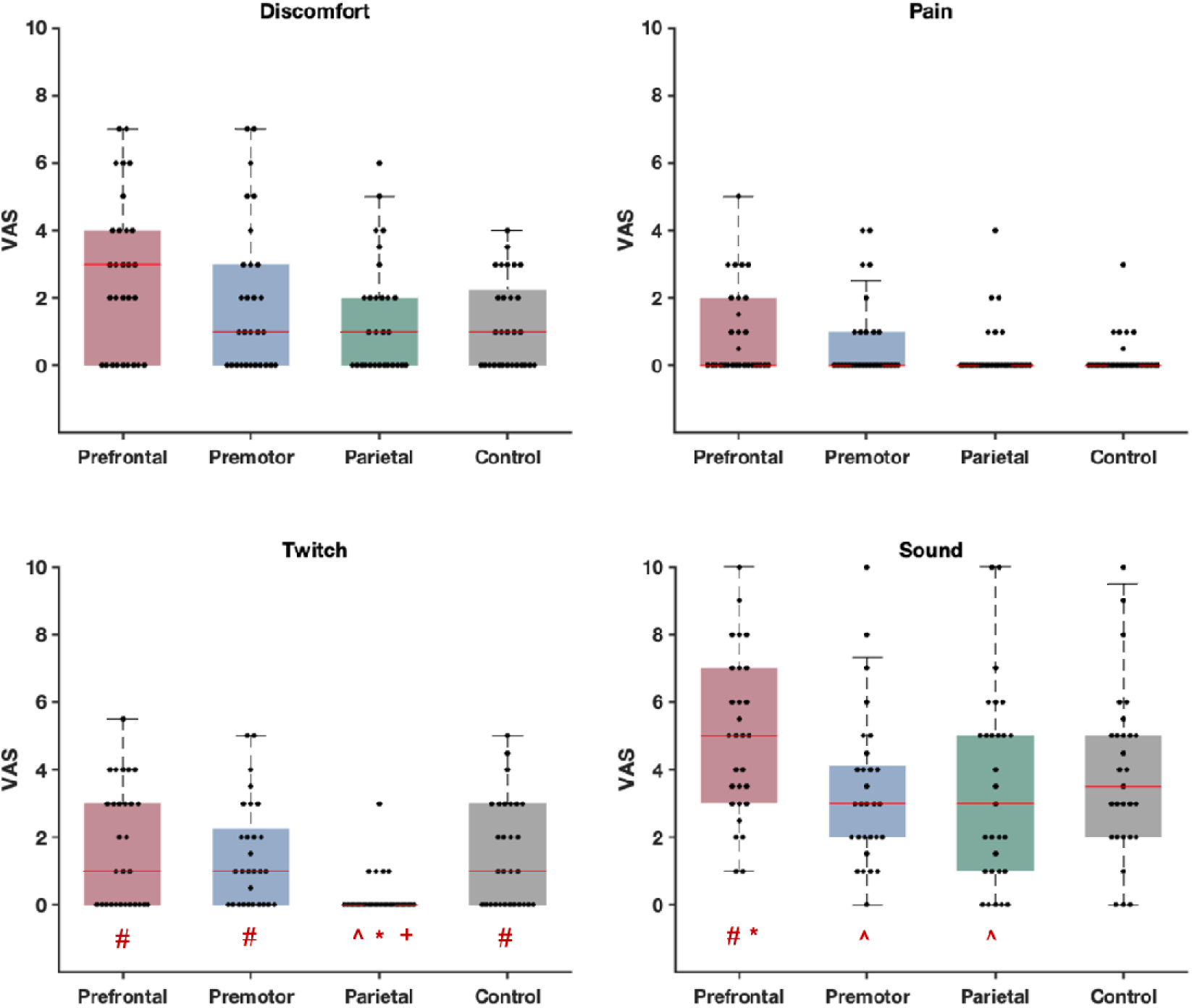
Experiment A-Self-reported perception of discomfort, pain, muscle twitch and click sound caused by real and control stimulation conditions. Each dot in the box and whisker plots represents the visual analogue scale (VAS) score for each individual. The shaded boxes highlight the 25th to 75th centiles of the values and the red horizontal lines within the boxes show the median of the values. **^**, **\***, **#,** and **+** indicate significant difference (FDR-corrected p<0.05) with prefrontal, premotor, parietal and control, respectively.

#### 3.1.2 Comparisons of TMS-evoked potentials following stimulation of different sites

Figure 3 illustrates the spatiotemporal distribution of TMS-evoked E-fields and TEPs following stimulation of each site. The TMS-evoked E-fields show clear spatial separation of the stimulated cortical regions when targeting the different cortical sites (Figure 3A). Together, the butterfly plots (Figure 3B), topoplots, and source estimations (Figure 3C) suggest differences in TEPs between stimulation sites within the first 60 ms following stimulation, which converge on a similar pattern between 60-300 ms between sites.

**Figure 3.**
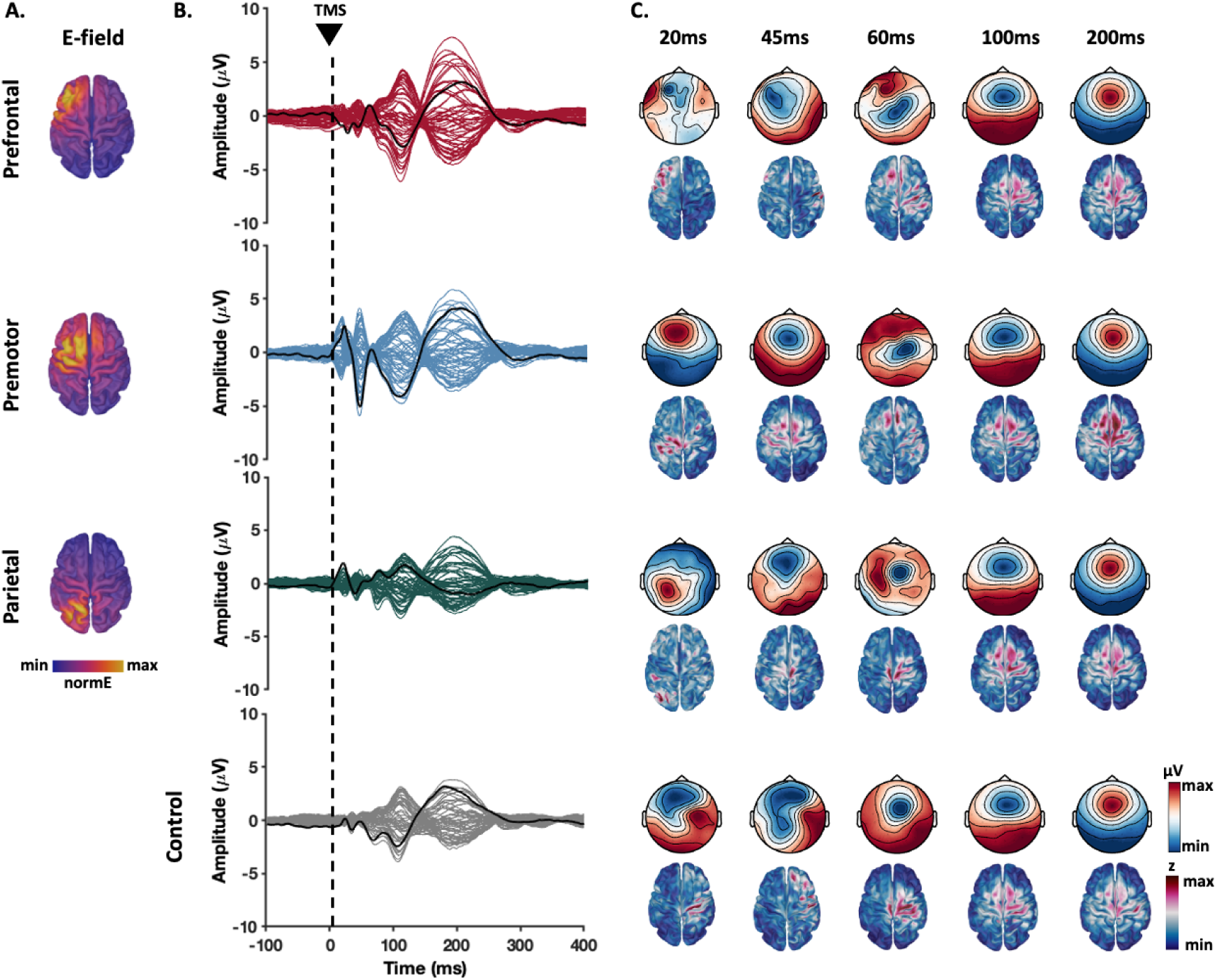
TMS-evoked potential and e-field distributions. A) Group average of the estimated e-fields in each stimulation condition. B) Butterfly plots indicate TEPs recorded by each electrode averaged across individuals and the thick black line represents the potentials recorded at the targeted site (prefrontal: F3; premotor: FC1, parietal: P3 and Shoulder: CZ). C) Topographical maps depict distribution of the potentials across scalp (upper maps) and their cortical source (lower maps) around the timepoints that TEP peaks appeared.

To compare similarities between TEPs following stimulation of different sites, we performed correlation analyses in both the temporal and spatial domain. Pairwise temporal correlations comparing the shape of TEPs within electrodes between sites were performed across two time windows, early (15-60 ms) and late (60-400 ms). The early time window revealed weak associations for most electrode comparisons, with the exception of a cluster of central electrodes which showed moderate strength correlations for most comparisons. In contrast, the late time window showed strong correlations between sites across all electrodes which peaked in frontocentral electrodes (Figure 4A), suggesting very similar TEP shapes between both active and control conditions. The pairwise assessments of spatial correlations (i.e., comparing the similarities of topoplots at each point in time) showed weak correlations between topographies within the first 60 ms post stimulation, with strong correlations present between 60-300ms post-stimulus peaking around 100 and 200 ms (Figure 4B). The strong temporal and spatial correlations from 60 ms between both active and control stimulation sites (e.g., the shoulder) suggest a common underlying neural source between conditions which is not dependent on direct cortical stimulation and is most likely the result of sensory input.

**Figure 4.**
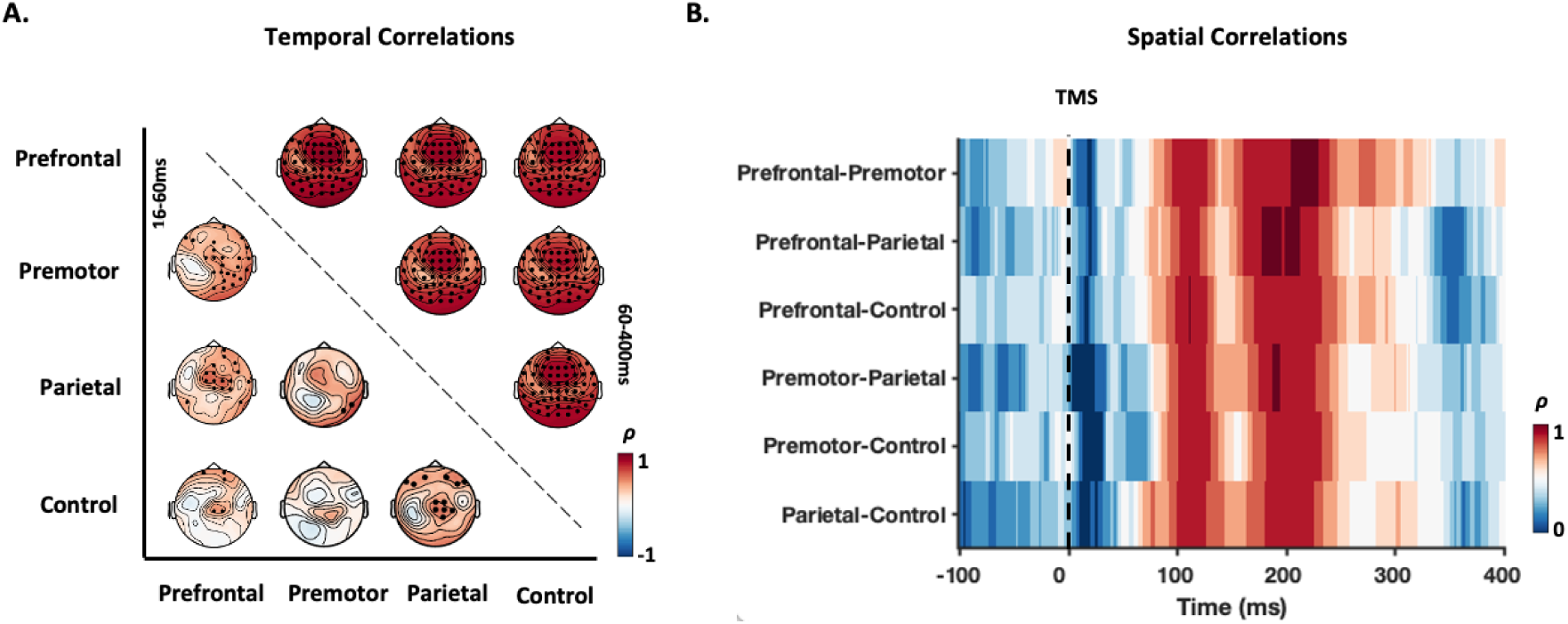
Pairwise spatial and temporal correlations between TEPs from different conditions. A) Spatial distribution of the Spearman’s correlation coefficients (ρ) between the potentials recorded by the same electrodes in each two conditions at two different time windows; early (16-60ms; lower triangle) and late (60-400ms; upper triangle). The channels highlighted in black demonstrated significant correlations (p _corrected_<0.05) between the two conditions. B) Temporal changes in Spearman’s correlation between the topographies of each two conditions.

To assess differences in the magnitude of TEPs between stimulation sites, we compared GMFP between sites across early (15-60 ms) and late (60-300 ms) time windows, corrected for differences in the peak E-field (Figure 5A,B). Comparisons of GMFP showed a significant difference between TEPs across conditions for early (one-way ANOVA: F (2, 84) = 5.32, p = 0.007) and late time windows (one-way ANOVA: F (2, 84) = 5.66, p = 0.005). Post hoc comparisons revealed that the premotor cortex showed larger GMFP compared to prefrontal (p = 0.04) and parietal (p = 0.01) cortex in the earlier time-window. In the late time window, prefrontal cortex GMFP was larger than parietal cortex (p = 0.006), showing a similar pattern to differences in sensory perceptions (Figure 2).

**Figure 5.**
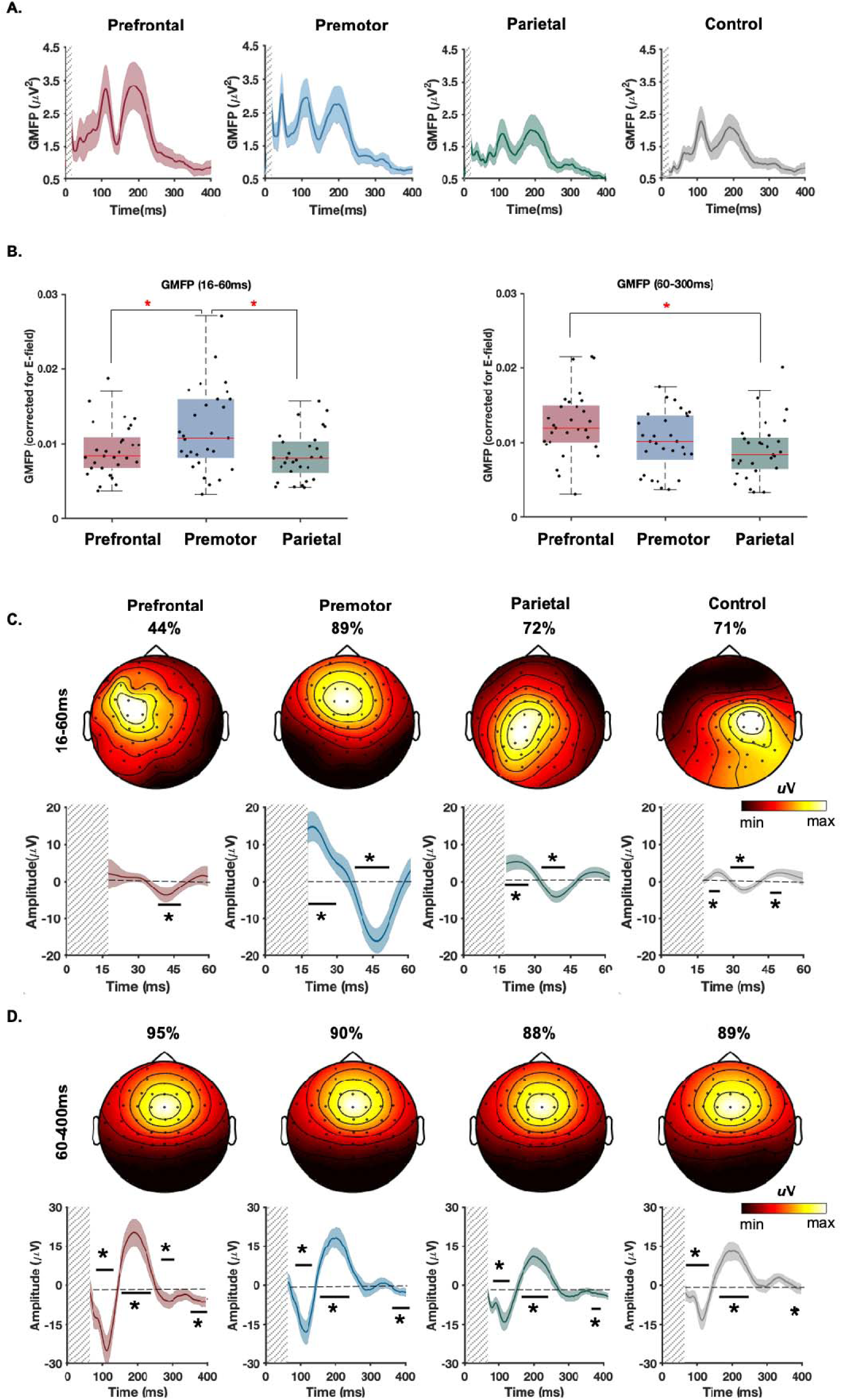
GMFP and the first principal component of the potentials recorded at each stimulation condition in experiment A. A) Fluctuations in GMFP following TMS. B) Comparison of GMFP across conditions at early and late time windows. Each dot in the box and whisker plots represents the value for each individual. The shaded boxes highlight the 25th to 75th centiles of the values and the red horizontal lines within the boxes show the median of the values. **\***indicates significant difference between the stimulation conditions (FDR-corrected p<0.05). C) The dominant PCs identified in early TEPs recorded between 16 and 60ms. D) The dominant PCs identified in late TEPs recorded between 60 and 400ms. The values above the scalp maps indicate the percentage of variance explained by the depicted component for each condition. The line graphs illustrate the changes of PCs amplitude over time. The thick lines represent the group averaged signal and the shaded areas show 95% CIs of the individual values. The vertical grey bars demonstrate the time-window of the potentials not considered for the analysis. The horizontal lines with * indicate when the signals significantly deviate from baseline (corrected p<0.05).

To further understand the similarities and differences between TEPs across the different time windows, we performed principal component analysis (PCA) on the early (16-60ms) and late (60-400ms) TEPs separately and identified the components most representative of the signals (i.e., explaining the maximum spatial variance) in the group-averaged data. TEPs from each individual were then weighted according to the detected PC maps to find the dominant temporal patterns. In line with the results from the correlation analyses, PCs from the earlier time-window demonstrated site-specific topographies, which tended to peak close to the site of stimulation (electrodes showing maxima for each condition; prefrontal = FC3; premotor = FCz; parietal = CP1). Moreover, the time series from PCs characterising the dominant early TEPs from each stimulated site showed differing peaks, including an N40 (peak = 42 ms) for prefrontal cortex, a P20 (peak = 19 ms) and N45 (peak = 46 ms) for premotor cortex, and a P20 (peak = 20 ms) and N35 (peak = 37 ms) for parietal cortex (Figure 5B). The peak-to-peak amplitude of the components was largest for the premotor; the condition with the strongest e-field (Figure 1C). In contrast, PCA of late TEPs revealed that at least 88% of the variance in the potentials were explained by similar frontocentrally distributed potentials for all sites (electrode showing maxima = FCz in all sites), with prominent N100 (maxima range = 111–115 ms across sites) and P200 peaks (maxima range = 191–202 ms across sites) (Figure 5C). Together, these findings suggest that early time periods are dominated by site specific TEPs which are consistent with activity resulting from stimulation of the targeted cortical region. In contrast, late time periods are dominated by TEPs with a common spatiotemporal pattern across stimulation sites including from non-cortical areas (i.e, the shoulder) which likely result from sensory input associated with the TMS pulse.

### 3.2 Experiment B

The first aim of experiment B was to assess whether the findings from experiment A replicated in a larger independent data set (n = 94) with different stimulation settings, in this case a lower relative TMS intensity (100% of rMT). TEPs are still present at lower TMS intensities (as low as 60% RMT (Komssi et al., 2004)) and threshold or subthreshold intensities are commonly used in TMS-EEG experiments to help minimise certain sensory inputs (Premoli et al., 2014). Stimulation parameters and the level of pulse perceptions for experiment B are presented in the supplementary materials (Table S1, Figure S1). As illustrated by Figure S2, the distribution of TEPs in experiment B closely resembled those observed in experiment A. All of the main findings replicated between studies, including: moderate temporal correlations between conditions in central electrodes during the early time period (Figure S2C); strong temporal and spatial correlations between conditions in the late time period (Figure S2D); site specific PC1s for the early time period (Figure S3A); and common frontocentral PC1s with N100 and P200 peaks for the late time period (Figure S3B). Note that e-field distribution was not estimated in experiment B, however the GMFP results resembled those from experiment A before correction, with higher GMFP amplitudes for the late time period corresponding to N100/P200 peaks in prefrontal compared with parietal cortex. Together, these findings show that the pattern of site-specific early TEPs and site-general late TEPs replicates for a lower TMS intensity, providing further evidence that the late TEPs likely reflect sensory input regardless of stimulation intensity.

The between-site comparisons from experiments A and B both show that later TEPs are larger in the prefrontal compared with the parietal cortex, a pattern which is also observed in self-report sensations of discomfort, scalp muscle twitch strength and click loudness. To investigate the contribution of different sources of sensory input to the amplitude of late TEPs, we took advantage of the large sample size in experiment B and stratified the groups based on the presence/absence of each sensory perception. Stratification was based on low perception (VAS < median) versus high level of perception (VAS > median) for the four examined sensations including sound (median VAS = 2), twitch (median VAS = 0), discomfort (median VAS = 1) and pain (median VAS = 0). When median was 0, all the individuals with no perception were assigned to the low VAS group. As there was no evidence for site specificity in the contribution of later potentials in our findings, we combined the data from all the three real stimulation conditions to further improve the power of analysis (both GMFP and the first PC). Wilcoxon signed-rank tests showed that late TEPs (>∼60ms) were mostly larger in the individuals with stronger perceptions. The difference in TEPs was prominent between the groups separated by sound perception across a wide range of time windows including the N100 and P200 peaks (Figure 6A). However, for muscle twitch and discomfort differences were mainly limited to earlier (∼80-90 ms) and later (∼220-280 ms) time periods. The results did not show a significant impact of pain perception on the magnitude of TEPs. This finding suggests that although both auditory and somatosensory inputs alter the amplitude of frontocentral signals, auditory inputs make a larger contribution to differences in the N100/P200 peaks. The low impact of somatosensory potentials observed in our findings could have potentially been driven by the confounding effect of different levels of noise masking across groups. As there were no correlations between VAS of sound and other sensations (all FDR-corrected p>0.05), each somatosensory group could have a mixture of both high and low levels of auditory perceptions. To make sure the sound masking was equivalent across the somatosensory comparisons, we selected a subset of participants reporting low VAS (<2) for sound perception, and stratified the groups based on the presence/absence of each somatosensory perception. The results did not reveal significant differences between groups with the exception of two brief windows for muscle twitch between 250 and 400 ms post TMS, suggesting that SEPs from the evaluated perceptions do make a small contribution to TEPs, but not necessarily at the N100/P200 peaks (Figure 6B).

**Figure 6.**
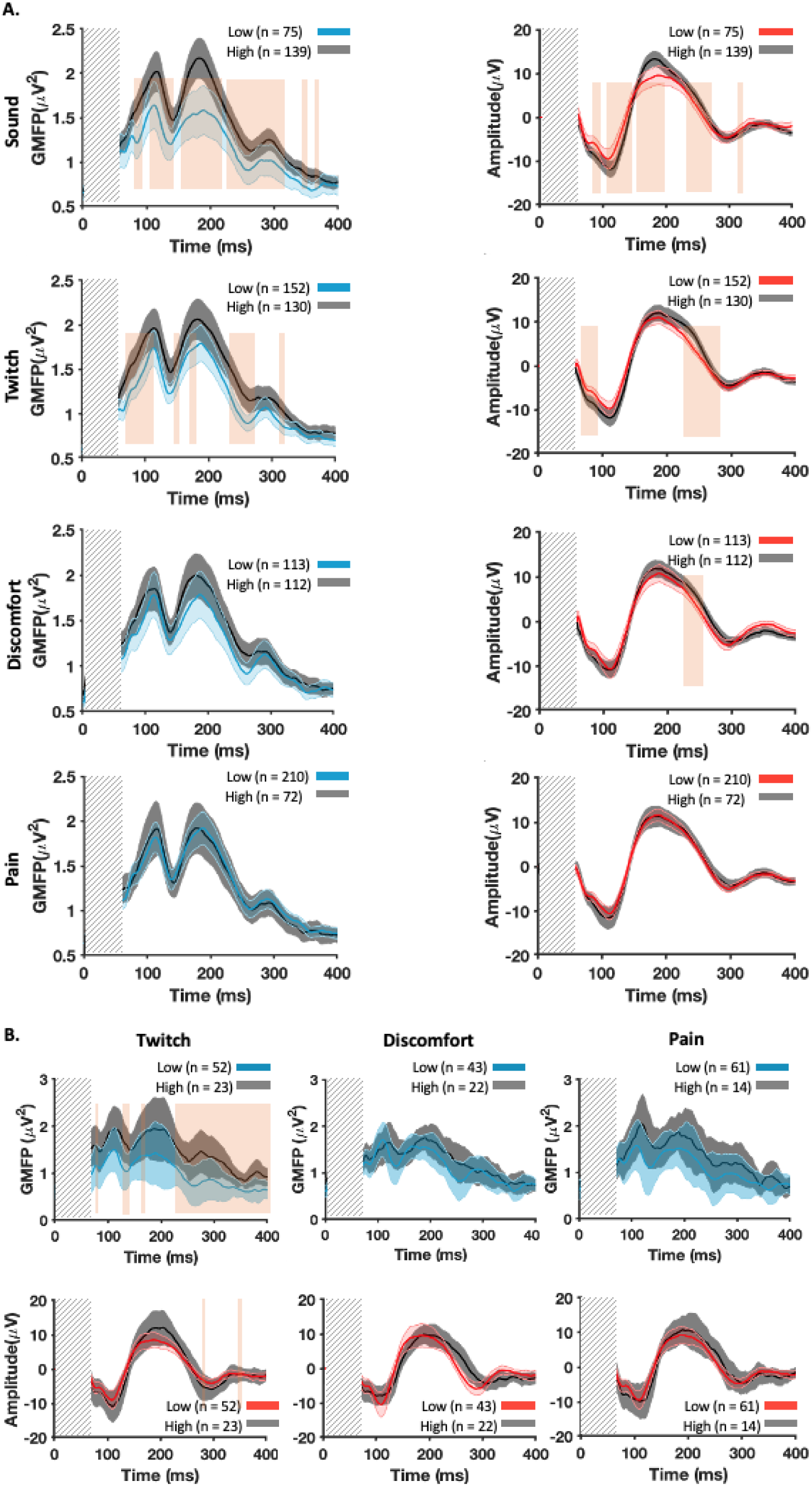
Experiment B-Comparisons of the late potentials between the individuals with high and low levels of sensory perceptions. A) Differences in the GMFP (left column) and the amplitude of the first PCs (left column) between the individuals who reported a low versus high level of sensory perception. B) Changes in the GMFP (upper row) and amplitude of the first PCs (lower row) from the subset of participants who perceived a low level of click sound (VAS<2). Potentials from all stimulation conditions are pooled into each group. The thick lines represent the group averaged signal and the shaded areas show 95% CIs of the individual values. The vertical grey bars demonstrate the time-window of the potentials not considered for the analysis. The orange vertical boxes cover the windows of time showing significant differences between groups (FDR-corrected P<0.05)

### 3.3 Experiment C

Findings from experiment A and B show that perception of the TMS click sound contributes to the N100/P200 peak amplitudes, and both click perception and N100/P200 amplitude differ between stimulation sites. To more directly assess the role of TMS sound perception in N100/P200 peak amplitude across stimulation sites, we performed an additional experiment which included similar conditions to experiment A, and a condition in which the auditory masking was removed altogether for each stimulation site while other sensations (i.e., discomfort, pain and muscle twitch) were kept at the same level (Figure 7A). We reasoned that if late, but not early TEPs contained AEPs, then increasing the amount of auditory input perceived by the participant should differentially alter the amplitude of early versus late TEPs. Furthermore, given that auditory masking was more successful in some sites (e.g., parietal cortex) compared to others (e.g., prefrontal cortex), we reasoned that late latency TEP peaks should change more when altering the level of auditory masking at parietal compared to prefrontal sites. TEPs from experiment C mostly replicated those from experiments A and B (table S2, figures S4-7). Similar to experiment A and B, auditory masking was more successful at reducing perception of the click sound for parietal cortex stimulation than prefrontal cortex stimulation (p = 0.012; Figure 7A). Removing the auditory masking increased the N100 and P200 for the parietal stimulation site (N100, p = 0.0004; P200, p = 0.0009), increased the P200, but not the N100 for the premotor stimulation site (N100, p = 0.12; P200, p = 0.021), and did not alter either peak for the prefrontal stimulation site (N100, p = 0.42; P200, p = 0.052). Earlier latency TEPs were not altered following the removal of auditory masking for any sites (all p>0.05). (Figure 7B). When comparing across stimulation sites, the change in late TEPs from no auditory masking to masking was significantly smaller in the prefrontal cortex compared to the premotor (N100, p = 0.01; P200, p = 0.07) and parietal (N100, p = 0.001; P200, p = 0.009) cortices (Figure 7C). This finding aligns with the variation observed in the efficiency of noise masking across sites, where participants perceived a higher level of click sound when stimulating the prefrontal cortex compared to the premotor (p = 0.04) and parietal (p = 0.03) cortices (Figure 7C). Together, these findings suggest: 1) auditory evoked potentials contribute to late, but not early latency TEPs, and 2) differences in the amplitude of late TEPs between stimulation sites is largely driven by differences in the success of auditory masking between sites.

**Figure 7.**
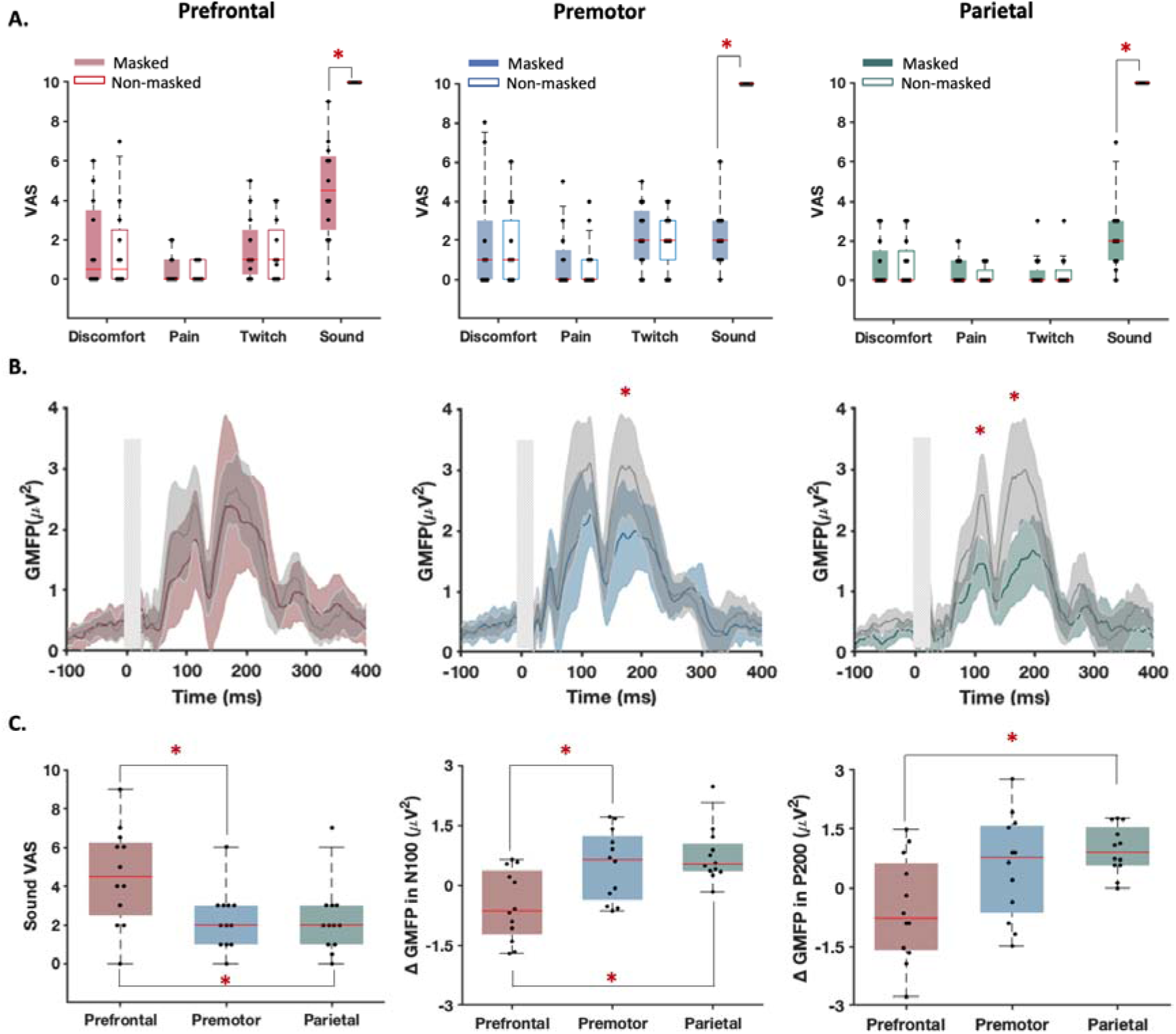
Experiment C: Sensory perceptions and TEPs from real (TMS with noise masking) and auditory control (TMS without noise masking) stimulation conditions. A) Self-reported perception of discomfort, pain, muscle twitch and click sound caused by each stimulation condition. B) Group-averaged GMFP evoked by TMS with and without noise masking. *indicates the significant difference (P<0.05) between conditions at the corresponding peak. The exact time-point for each peak was defined as when the group averaged GMFP from the real TMS condition had the largest value within the range of ±15ms around the specified time (e.g., between 85 and 115ms for N100). The thick line represents the group-averaged values. The shaded areas show 95% CIs of the values and the vertical grey bars demonstrate the time-window that was not considered for the analysis. C) Comparing the stimulation sites for the efficiency of noise masking and GMFP changes at N100 (100ms) and P200 (200ms) following removal of noise masking. A, C) Each dot in the box and whisker plots represents the visual analogue scale (VAS) score for each individual. The shaded boxes highlight the 25th to 75th centiles of the values and the red horizontal lines within the boxes show the median of the values. **\***indicates significant difference between the stimulation conditions (FDR-corrected p <0.05).

An additional limitation from experiments A and B is that the sensory control condition was performed by stimulating the shoulder, which does not control for spatial differences in somatosensory input when stimulating different cortical/scalp locations. To address this limitation, we performed a further site-specific control condition which involved applying a weak electrical stimulation to the same scalp location while the TMS coil was held away from the scalp, thereby preventing active cortical stimulation by the TMS pulse. The perceived sensations were comparable between the two conditions in all stimulation sites except that in the prefrontal cortex the click sound was louder in real stimulation (p = 0.009) (Figure 8A). Spatial correlations showed significant associations between the real and control TEPs from ∼60 ms, maximising around 100 and 200 ms; in accordance with the N100/P200 complex of AEPs (Figure 8B). As depicted in Figure 8B-C, this correlation pattern closely resembles that between TEPs and PEPs from shoulder stimulation further suggesting a minimal contribution from somatosensory inputs potentially triggered by scalp stimulation to early TEPs.

**Figure 8.**
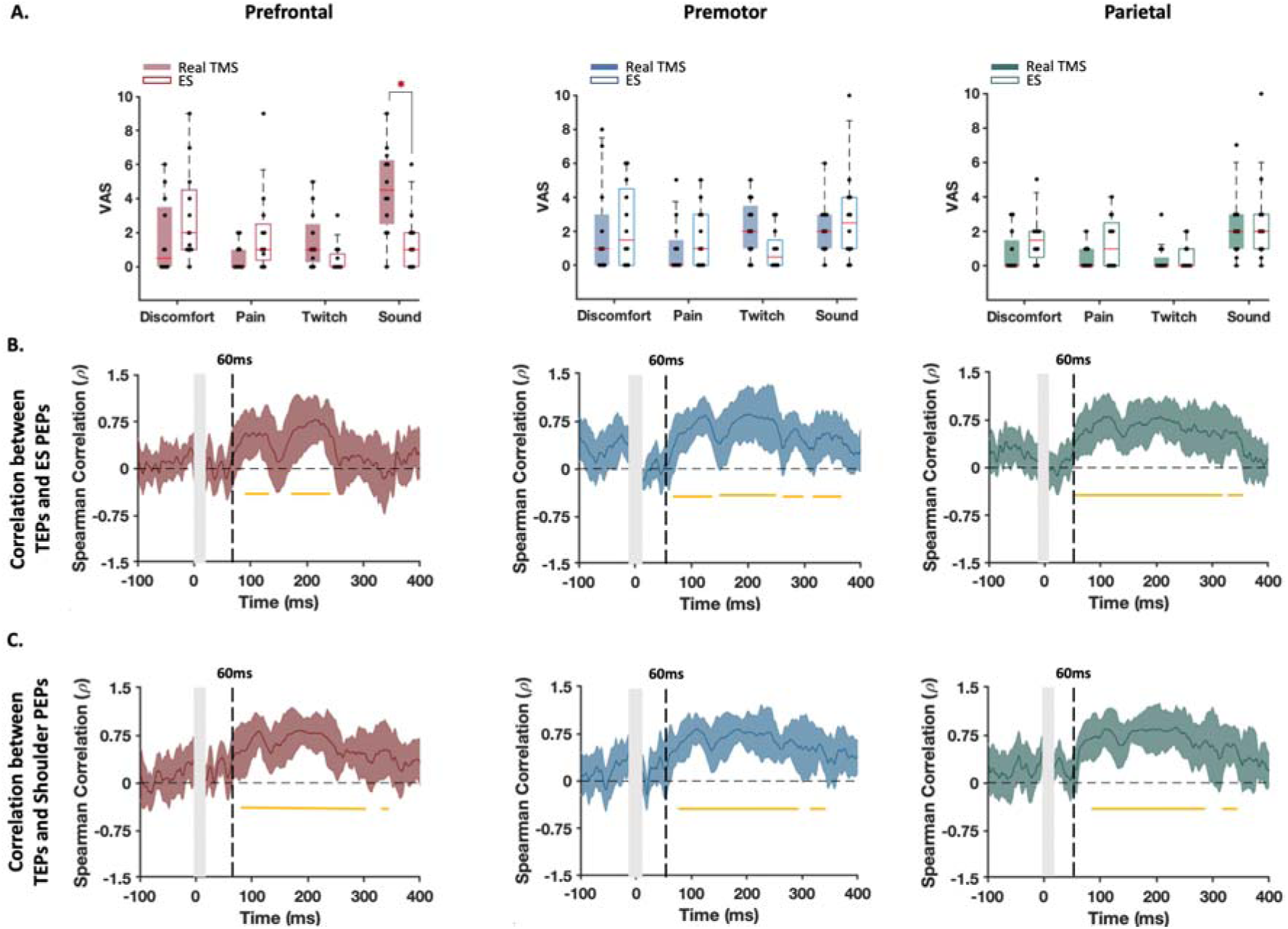
Experiment C: Sensory perceptions and TEPs from real TMS and somatosensory control (electrical stimulation) conditions. A) Self-reported perception of discomfort, pain, muscle twitch and click sound caused by each stimulation condition. Each dot in the box and whisker plots represents the visual analogue scale (VAS) score for each individual. The shaded boxes highlight the 25th to 75th centiles of the values and the red horizontal lines within the boxes show the median of the values. **\***indicates significant difference between the stimulation conditions (FDR-corrected p <0.05). B-C) Changes in correlations between the GMFP of the responses to real and sham stimulations over time. The horizontal lines below the zero line indicate the windows of significant correlations. The thick line represents the group-averaged values. The shaded areas show 95% CIs of the values and the vertical grey bars demonstrate the time-window that was not considered for the analysis.

## 4. Discussion

In the present study, we examined the EEG responses to TMS following stimulation of three non-motor regions (prefrontal, premotor and parietal cortex) and evaluated their associations with the EEG responses to sensory stimulation. We aimed to identify the features in TEPs that were sensitive to auditory and somatosensory inputs and those which were not. There were four main findings. First, we replicated the strong temporal and spatial correlations between TEPs from different sites and sensory evoked potentials between 60-300 ms, which peaked around the frontocentral N100 and P200 potentials, suggesting these late peaks primarily reflect sensory potentials. We showed that these relationships are replicated for different stimulation intensities (100% and 120% of scalp-to-cortex adjusted RMT) and for different types of sensory stimulation (shoulder stimulation and site-specific electrical stimulation of the scalp with a coil spacer). Second, we found that sensory experiences differed following stimulation of different sites, with prefrontal cortex stimulation resulting in more discomfort, stronger scalp muscle twitches, and louder perception of the coil click (despite white noise masking) compared to the other sites. We also found that the N100/P200 peaks were larger in the prefrontal cortex compared to the parietal cortex, showing a similar pattern to sensory experience. Third, we observed that late TEPs were stronger in individuals who perceived louder click sounds and had larger muscle twitches, with stronger click sound perception in particular resulting in a larger N100/P200 peak amplitude. Fourth, we found that removing noise-masking increased N100/P200 amplitude to differing extents across different stimulation sites, with both click perception and TEP amplitudes more strongly modulated at parietal than prefrontal cortex. Importantly, across all experiments the early latency TEPs (<60 ms) showed site-specific topographies and time courses and were not altered following any sensory comparison or manipulation, suggesting that these peaks most likely reflect neural activity resulting from direct stimulation of the cortex by TMS. Together, these findings provide strong evidence that later TEP peaks are sensitive to auditory and somatosensory experiences of TMS, which can differ between stimulation sites across the scalp, whereas earlier peaks likely reflect activity initiated by transcranial stimulation of the cortex.

A growing number of studies have shown strong correlations between frontocentral N100 and P200 PEPs resulting from sensory sham conditions and TEPs following stimulation of multiple sites, including motor, prefrontal, premotor, parietal and visual cortex (Biabani et al., 2019; Chowdhury et al., 2022; Conde et al., 2019; Freedberg et al., 2020; Gordon et al., 2021; Herring et al., 2015; Rocchi et al., 2021). The present findings replicate this relationship in prefrontal, premotor, and parietal cortex across three independent datasets (total n=135), providing further evidence that PEPs contribute to TEPs regardless of stimulation site, stimulation intensity, and the method of sensory stimulation used for comparison (e.g., shoulder vs electrical scalp stimulation). While we were not able to replicate the strong spatial correlations between PEPs and prefrontal/parietal TEPs at ∼25 ms and ∼40 ms as reported by Conde, Tomasevic and colleagues (Conde et al., 2019), we did see a moderate temporal correlation in central electrodes between conditions during early (16-60 ms) time windows. However, we also observed site specific TEPs with independent spatial and temporal profiles which dominated the early time windows, suggesting some early peaks are primarily reflective of transcranial evoked activity. We replicated previous studies showing the strength of self-reported sensory experiences such as discomfort, scalp muscle twitches, and perceived coil clicks (regardless of masking) differed across TMS scalp locations (Meteyard and Holmes, 2018), and were larger/stronger following stimulation of prefrontal compared to parietal cortex. We extended these findings by showing the stimulation sites with stronger sensory experience were accompanied by larger N100/P200 peak amplitudes, a relationship which was not observed in early latency TEPs. These findings provide an important insight suggesting that the contribution of PEPs to TEPs may differ across stimulation sites depending on differing sensory experiences.

An important unresolved question is the relative contribution of auditory and somatosensory experience to PEPs following TMS. Some studies have argued PEPs are entirely auditory in origin (Ilmoniemi and Kičić, 2010; Paus et al., 2001; Rocchi et al., 2021), whereas others have suggested both auditory and somatosensory input contributes to PEPs (Conde et al., 2019; Gordon et al., 2021). We used between subject comparisons to isolate the impact of different sensory perceptions including click sound, muscle twitch, pain and discomfort on the magnitude of the later responses to TMS. We found that TEPs were significantly larger in individuals with stronger perception of the TMS click sound and TMS-evoked scalp muscle twitches, with stronger perception of sound in particular resulting in larger frontocentral N100/P200 peak amplitudes. In a follow-up experiment, we found that removing auditory masking increased frontocentral N100/P200 peak amplitudes within participants without altering early latency TEPs, replicating similar findings from the motor cortex (Rocchi et al., 2021; Ter Braack et al., 2015). Furthermore, the modulation of TEP amplitudes was stronger in the sites with more effective auditory masking. Together, these results suggest that auditory input is the major contributor to PEPs in TEPs, although somatosensation also makes a contribution. In line with our findings, the N100/P200 complex is observed following a range of different sensory stimuli (Mouraux and Iannetti, 2009; Singhal et al., 2002), suggesting this ERP may reflect higher-level processing of perceptual inputs from a broad cortical network (Downar et al., 2002). The size and distribution of N100/P200 potentials makes it possible that other TEPs were masked by the strong PEPs (Gordon et al., 2023; Ross et al., 2022a). Both online and offline methods for minimising PEPs may help uncover these additional TEPs. In addition to a generalised sensory response, we also observed early latency TEPs (<60 ms) that were specific to the site of stimulation, with peaks largest in the electrodes near the stimulated cortical region. This finding replicates a growing number of studies that have reported site-specificity of early TEPs (Casarotto et al., 2010; Fecchio et al., 2017; Kähkönen et al., 2004; Ozdemir et al., 2020; Passera et al., 2022; Rogasch et al., 2020; Rosanova et al., 2009). More specifically, we replicated TEP peaks including a N40 following dorsolateral prefrontal cortex stimulation (Eshel et al., 2020; Kerwin et al., 2018; Rogasch et al., 2015), a P20 and N45 following premotor cortex stimulation (Belardinelli et al., 2019; Casarotto et al., 2022, 2011; Rosanova et al., 2009), and a P20 and N35 following parietal cortex stimulation (Belardinelli et al., 2019; Casarotto et al., 2010; Rosanova et al., 2009). While the origin of these early TEPs is unclear, several lines of evidence suggest these peaks likely reflect neural activity generated by the site of stimulation. First, as observed in this study, site-specific early TEPs show weak correlations with PEPs from sensory control conditions and show spatial characteristics consistent with activity from the site of stimulation. Second, early TEPs are not altered by changing sensory experiences of TMS, like the level of TMS click perception. Third, early TEPs are only present when healthy cortical tissue is stimulated, not when lesioned tissue is stimulated (Gosseries et al., 2015). Finally, site specific early TEPs with similar latencies and invariance to sensory input have also been observed from invasive cortical recordings in non-human primates (Perera et al., 2023). Together, these findings suggest that early latency TEPs most likely reflect neural activity resulting from transcranial stimulation of the cortex.

### 4.1 Limitations

There are several important limitations to the current study. First, we used white noise played through in-ear headphones to mask the clicking sound of TMS. Currently, there are two main approaches for masking the TMS clicking sound: playing white noise through headphones or playing noise with a frequency spectrum adapted from the TMS click sound itself (Russo et al., 2022). While certain studies have suggested that the two are roughly equivalent in reducing the N100/P200 amplitude (Ter Braack et al., 2015), more recent studies have shown that adapted noise is more effective at reducing the participants perception of the clicking sound than white noise masking (Russo et al., 2022). Indeed, near complete suppression of the N100/P200 peak has been reported when using adapted noise masking across several stimulation sites (Massimini et al., 2005; Rocchi et al., 2021). The subjective ratings of click sound perception in our study suggested that white noise was not able to completely mask the TMS click sound for most individuals and conditions. It is possible that more effective suppression of the TMS click sound perception could have been achieved if adapted noise masking was used instead of white noise in the current studies. However, even adapted noise masking does not appear effective for all stimulation sites. For example, a recent study showed no reduction in the N100/P200 peak amplitude between no masking and adapted noise masking following DLPFC stimulation (Ross et al., 2022b), similar to the results of our study. Second, while the site-specific characteristics of the early latency TEPs (i.e., <60 ms) are consistent with activity evoked by transcranial stimulation of the cortex, we cannot rule out that sensory potentials also contribute in some way to these early peaks. Changing the level of auditory input did not alter early TEP amplitudes for any of the stimulation sites, making the contribution of auditory potentials unlikely. However, the contribution of somatosensory potentials is less clear. Recently, a sham condition was developed which experimentally saturates somatosensory input by using concurrent electrical stimulation of the scalp with TMS (Gordon et al., 2021). This approach was used to rule out somatosensory contributions to early TEPs following motor cortex stimulation (Gordon et al., 2023). A similar approach may prove useful in further assessing the contribution of somatosensory potentials to early TEPs following stimulation of other cortical sites.

### 4.2 Implications

An important implication of the current findings is that both auditory and somatosensory input can induce potentials which contribute to TEPs. While it is possible that auditory input can be minimised across different sites with adequate noise masking, controlling the level of somatosensory input is much more challenging, especially for more lateral sites which experience higher levels of TMS-evoked muscle twitches like the DLPFC. Therefore, it may not be possible to completely eliminate the contribution of sensory potentials from TEPs using experimental arrangements for certain stimulation sites. As such, the use of well-designed experimental control conditions and the continued development of off-line approaches is required for assessing and minimising sensory potentials in TEPs (Hernandez-Pavon et al., 2023; Rogasch et al., 2022). Our findings also highlight the importance of considering sensory experiences when comparing TEPs between different stimulation sites and different groups which may have differences in sensory perception. For example, older individuals (Kok, 2000) and individuals with psychiatric (Javitt, 2009; Shen et al., 2020) and neurological (Horvath et al., 2018) conditions show differences in sensory evoked potentials. Therefore, differences in TEPs between these groups could be attributed to either changes in sensory perception or the physiology of targeted neural circuits following TMS or a combination of the two. Careful experimental design is required to disentangle these possible contributions in between-group TMS-EEG studies.

### 4.3 Conclusions

The present findings suggest that short-latency TEPs (∼16-60ms) most likely represent neural activity resulting from transcranial stimulation of the cortex as they were: 1) lateralized to the site of stimulation; 2) uncorrelated with sensory evoked potentials from sensory control conditions without active TMS; and 3) invariant to changes in the level of TMS click sound perception. In contrast, frontocentral TEPs particularly around the N100/P200 peaks most likely reflect sensory evoked potentials as they: 1) showed high spatial and temporal correlations between stimulation sites and with sensory control conditions; 2) differed in amplitude between individuals who experienced more vs less sensory input from TMS; and 3) differed in amplitude within individuals when the level of TMS click sound perception was modulated. Importantly, we also showed that later TEP amplitudes were larger following stimulation of sites with more sensory input like the DLPFC and were larger in individuals with higher levels of click sound perception and TMS-evoked scalp muscle twitches, suggesting both auditory and somatosensory input contributes to TEPs. Our findings have important implications for the design and interpretation of TMS-EEG studies, suggesting sensory potentials may be more difficult to control experimentally for certain stimulation sites where muscle twitches are unavoidable. Furthermore, our findings highlight the importance of accounting for sensory potentials when comparing TEPs between stimulation sites and participant cohorts with different levels of sensory perception.

### 5.1 Data and code availability

Processed TMS-EEG data from experiments A-C are available at https://doi.org/10.26180/24514741. Analysis code is available at: https://github.com/ManaBiabani/NonMotor_TEPs_PEPs.git.

### 5.2 Author contributions

MB, AF and NCR conceptualised and designed the experiments. Data collection was carried out by MB, ST, LG. Data analysis was performed by MB and NR who also took the lead in interpreting the results and drafting the manuscript. MG, JGS, GO, MAB and AF provided critical reviews. All the authors contributed to the interpretation of the results, edited, and approved the manuscript.

### 5.3 Funding

This research work was supported by the Australian Research Council, Australia [DP170100738, DE180100741, FT210100694].

## Supporting information

Supplemental Results

