## Supplemental Results for "Characterising the contribution of auditory and somatosensory inputs to TMS-evoked potentials following stimulation of prefrontal, premotor and parietal cortex"

**Table S1** Mean±SD of stimulation parameters (Experiment B)

| | Distance (mm) | Intensity (%MSO) |
|---|---|---|
| **Prefrontal** | 18.15±3.83 | 56.59±12.12 |
| **Premotor** | 21.30±4.57 | 62.02±16.94 |
| **Parietal** | 20.02±4.53 | 61.67±12.59 |
| **Shoulder** | – | 52.77±10.73 |

Distance: scalp-to-cortex distance; Intensity: Distance-adjusted stimulation intensity; %MSO: Percentage of maximum stimulator output.

**Table S2** Mean±SD of stimulation parameters (Experiment C)

| | Distance (mm) | TMS Intensity (%MSO) | ES Intensity (mA) |
|---|---|---|---|
| **Prefrontal** | 18.71±4.44 | 38.5±11.02 | 0.65±0.16 |
| **Premotor** | 21.4±6.68 | 44.42±7.64 | 0.73±0.31 |
| **Parietal** | 23.01±6.18 | 46.92±8.22 | 0.73±0.30 |
| **Shoulder** | – | 45.52±10.83 | – |

Distance: scalp-to-cortex distance; Intensity: Distance-adjusted stimulation intensity; %MSO: Percentage of maximum stimulator output.

Figure 1s and 2s illustrates the levels of perception of pulses across the stimulated regions in experiments B. According to Shapiro-Wilk normality tests, VAS values were not normally distributed in all conditions (p>0.05). Therefore, we used Kruskal-Wallis tests for comparisons across all conditions, followed by FDR corrected Wilcoxon Signed-Rank tests for pairwise comparisons. Despite the differences in stimulation intensities, participants in experiment B showed almost the same pattern of perception across cortical sites as observed in experiment A; with prefrontal and Parietal causing the strongest and weakest sensations, respectively. While pain was perceived comparably across stimulation conditions (Chi-square = 6.82, df = 3, p = 0.077) the other sensations showed significant differences (For discomfort Chi-square = 25.76, df = 3, p <0.0001; Pairwise FDR-Corrected p : prefrontal-premotor = 0.0005, prefrontal-Parietal <0.0001, prefrontal-control<0.0001, premotor-Parietal = 0.04, premotor-control = 0.16; Parietal-control = 0.92; For twitch Chi-square = 28.73, df = 3, p <0.0001; Pairwise FDR-Corrected p : prefrontal-premotor <0.0001, prefrontal-Parietal <0.0001, prefrontal-control = 0.0003, premotor-Parietal = 0.04, premotor-control = 0.24 and Parietal-control = 0.009; For sound Chi-square = 19.69, df = 3, p = 0.0002; Pairwise FDR-Corrected p : prefrontal-premotor = 0.04, prefrontal-Parietal <0.0001, prefrontal-control<0.0001, premotor-Parietal = 0.0005, premotor-control = 0.001, Parietal-control = 0.34) (Figure 1s).

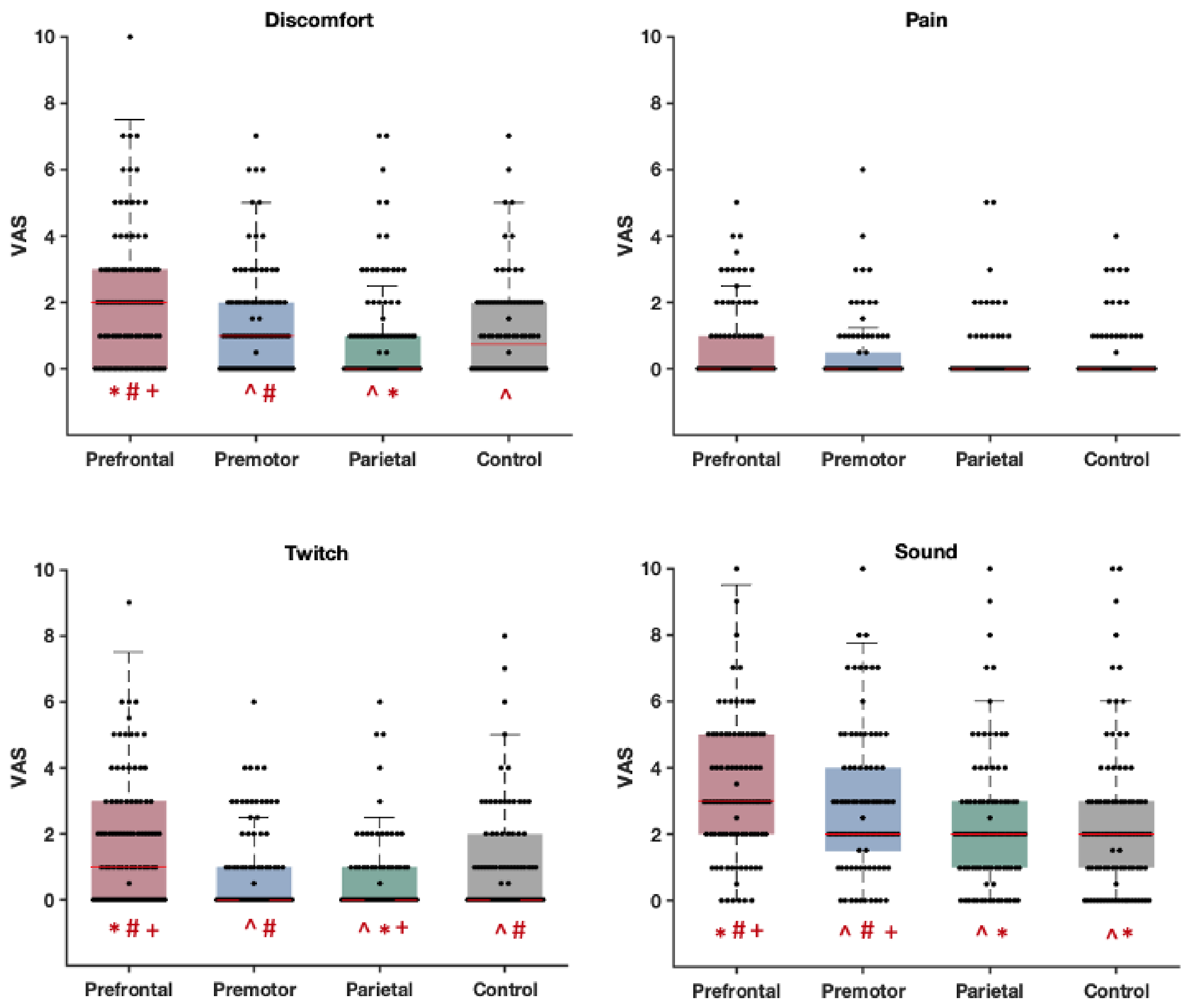


**Figure S1. Experiment B- Self-reported perception of discomfort, pain, muscle twitch and click sound caused by real and control stimulation conditions**. Each dot in the box and whisker plots represents the visual analogue scale (VAS) score for each individual. The shaded boxes highlight the 25th to 75th centiles of the values and the red horizontal lines within the boxes show the median of the values. ^, *****, #, and + indicate significant difference (FDR-corrected $p<0.05$) with prefrontal, premotor, Parietal and control, respectively.

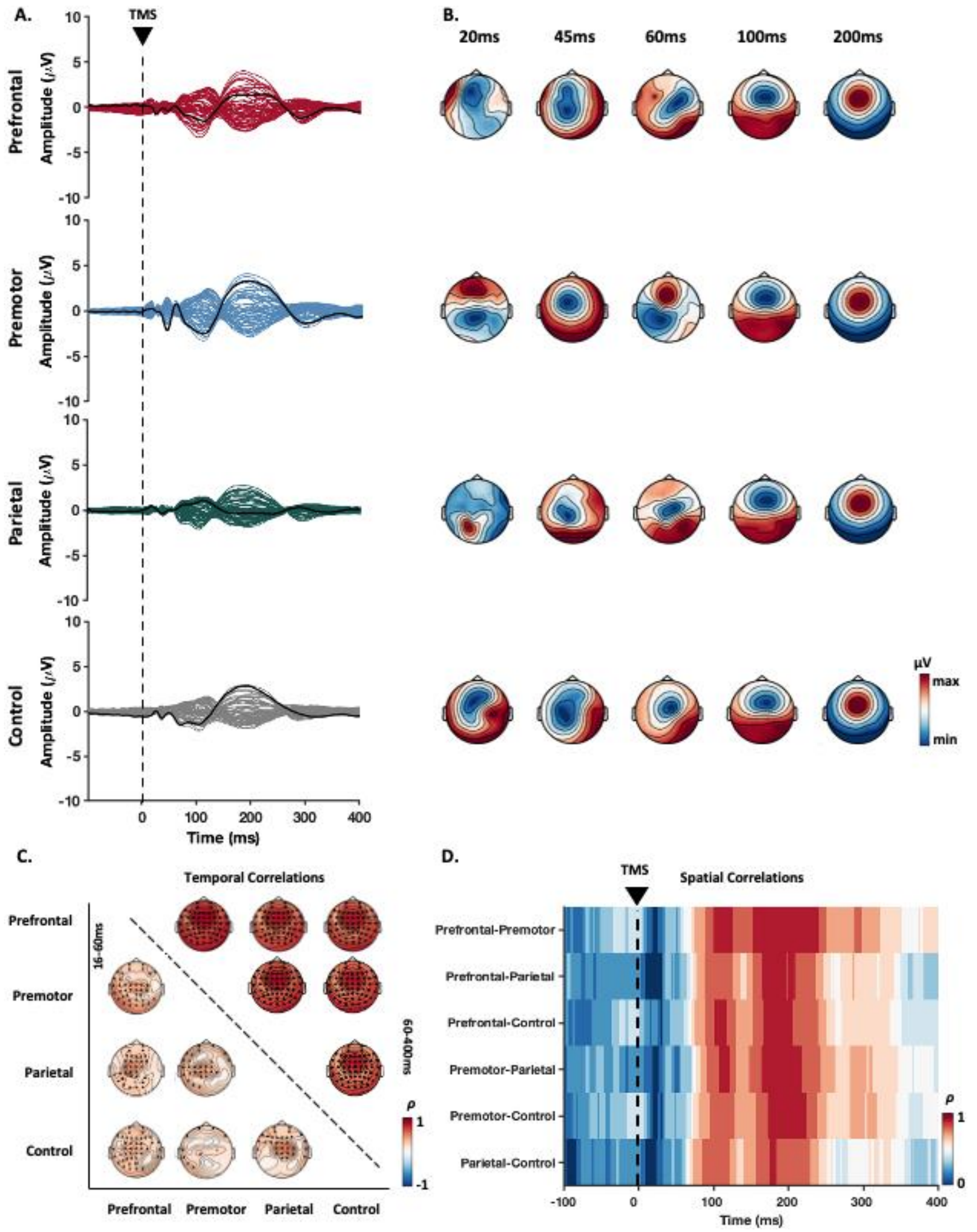


**Figure S2. Experiment B- using subthreshold TMS: Spatiotemporal distribution of TEPs evoked by TMS over four different areas and their correlation among conditions.**

A) Butterfly plots indicate TEPs recorded by each electrode averaged across individuals and the thick black line represent the potentials recorded at the targeted site (prefrontal: F3; premotor: FC1, Parietal: P3 and Shoulder: CZ). B) Topographical maps depict distribution of the potentials across scalp around the timepoints that TEP peaks appeared. C) Spatial distribution of the Spearman's correlation coefficients (ρ) between the potentials recorded by the same electrodes in each two conditions at two different time windows; early (16-60ms; lower triangle) and late (60-400ms; upper triangle). The channels highlighted in black demonstrated significant correlations ($p_{corrected}<0.05$) between the two conditions. D) Temporal changes in Spearman's correlation between the topographies of each two conditions.

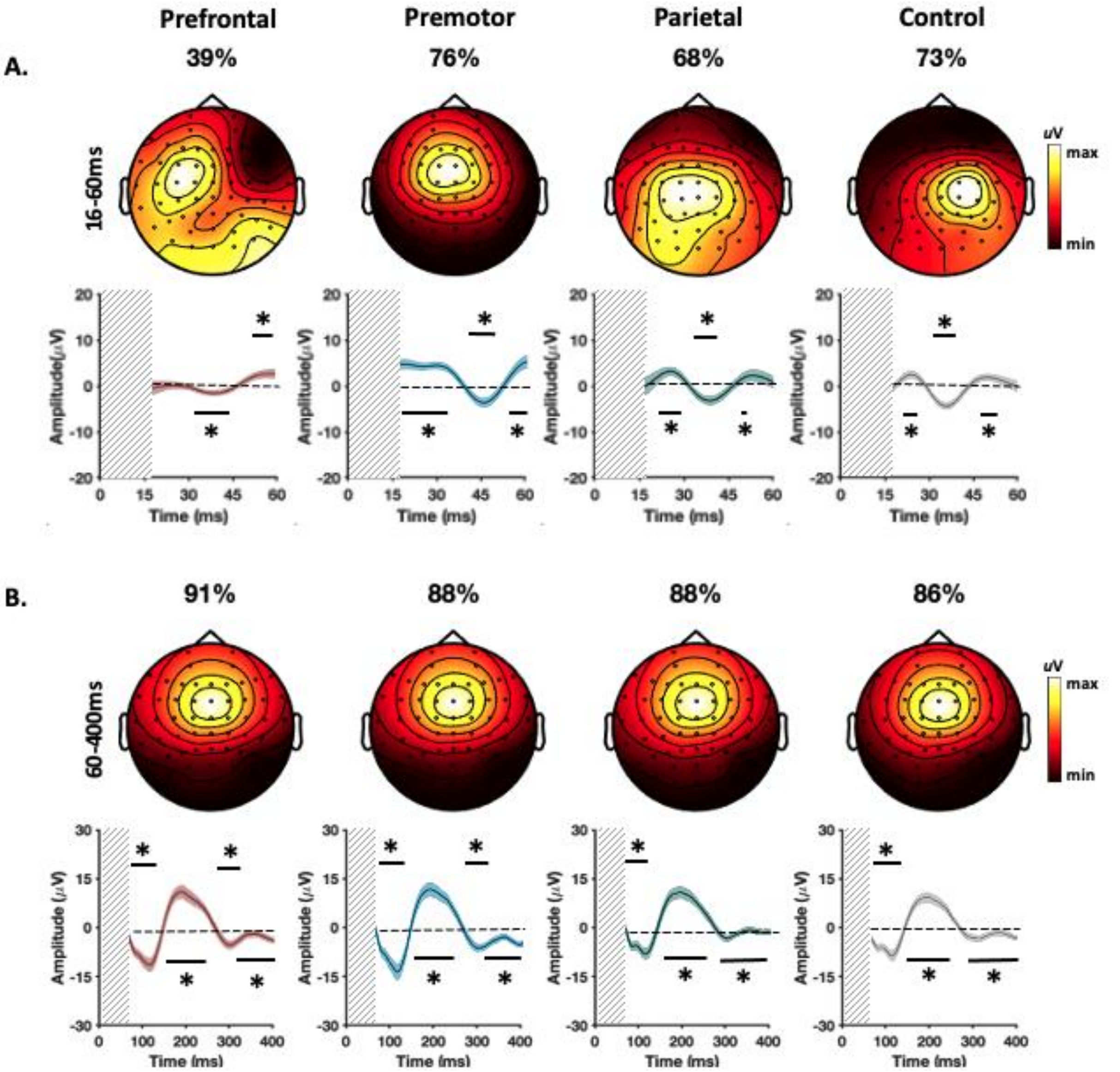


**Figure S3. Experiment B- The principal components explaining the maximum variance of the potentials recorded at each stimulation condition.** A) The dominant PCs identified in early TEPs recorded between 16 and 60ms. B) The dominant PCs identified in late TEPs recorded between 60 and 400ms. The values above the scalp maps indicate the percentage of variance explained by the depicted component for each condition. The line graphs illustrate the changes of PCs amplitude over time. The thick lines represent the group averaged signal and the shaded areas show 95% CIs of the individual values. The vertical grey bars demonstrate the time-window of the potentials not considered for the analysis. The horizontal lines with * indicate when the signals significantly deviate from baseline (corrected p<0.05).

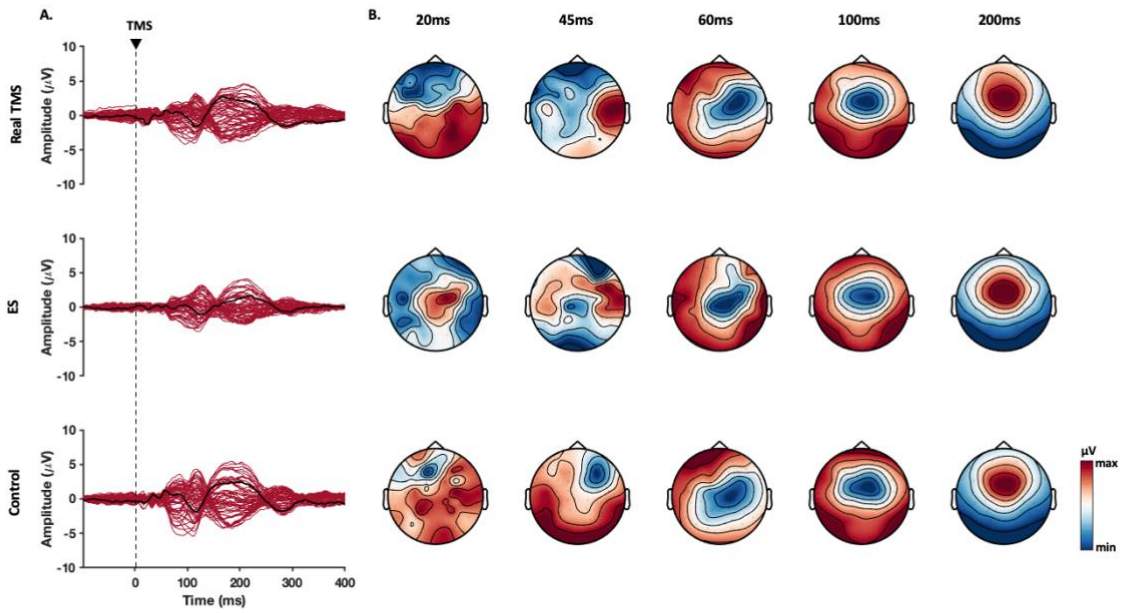


**Figure S4. Experiment C- Spatiotemporal distribution of scalp recorded EEG potentials evoked by different stimulation conditions over the prefrontal cortex.** In real TMS condition, stimulation was applied over the targeted area while the standard noise masking procedure was adopted. In the ES condition, electrical stimulation was applied over the targeted area, TMS coil was placed over the ES electrode tilted at 90° and the standard noise masking procedure was adopted. In control condition, TMS was applied without noise masking. B) Topographical maps depict distribution of the potentials across the scalp.

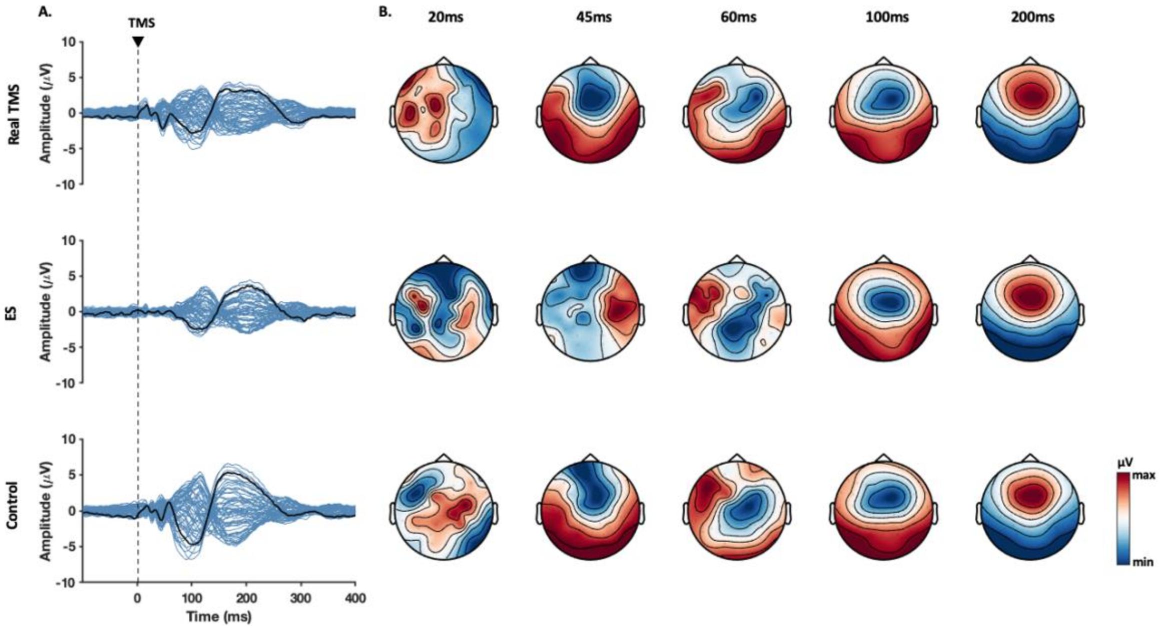


**Figure S5. Experiment C- Spatiotemporal distribution of scalp recorded EEG potentials evoked by different stimulation conditions over the premotor cortex.** In real TMS condition, stimulation was applied over the targeted area while the standard noise masking procedure was adopted. In the ES condition, electrical stimulation was applied over the targeted area, TMS coil was placed over the ES electrode tilted at 90° and the standard noise masking procedure was adopted. In control condition, TMS was applied without noise masking. B) Topographical maps depict distribution of the potentials across the scalp.

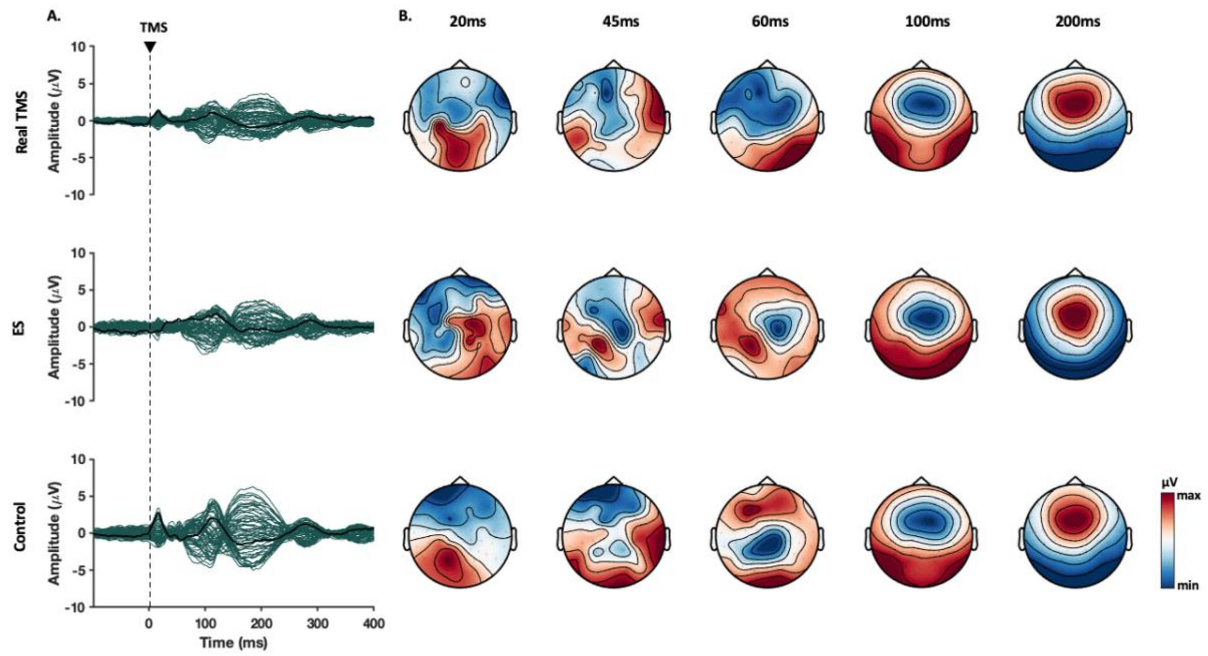


**Figure S6. Experiment C- Spatiotemporal distribution of scalp recorded EEG potentials evoked by different stimulation conditions over the parietal cortex.** In real TMS condition, stimulation was applied over the targeted area while the standard noise masking procedure was adopted. In the ES condition, electrical stimulation was applied over the targeted area, TMS coil was placed over the ES electrode tilted at 90° and the standard noise masking procedure was adopted. In control condition, TMS was applied without noise masking. B) Topographical maps depict distribution of the potentials across the scalp.

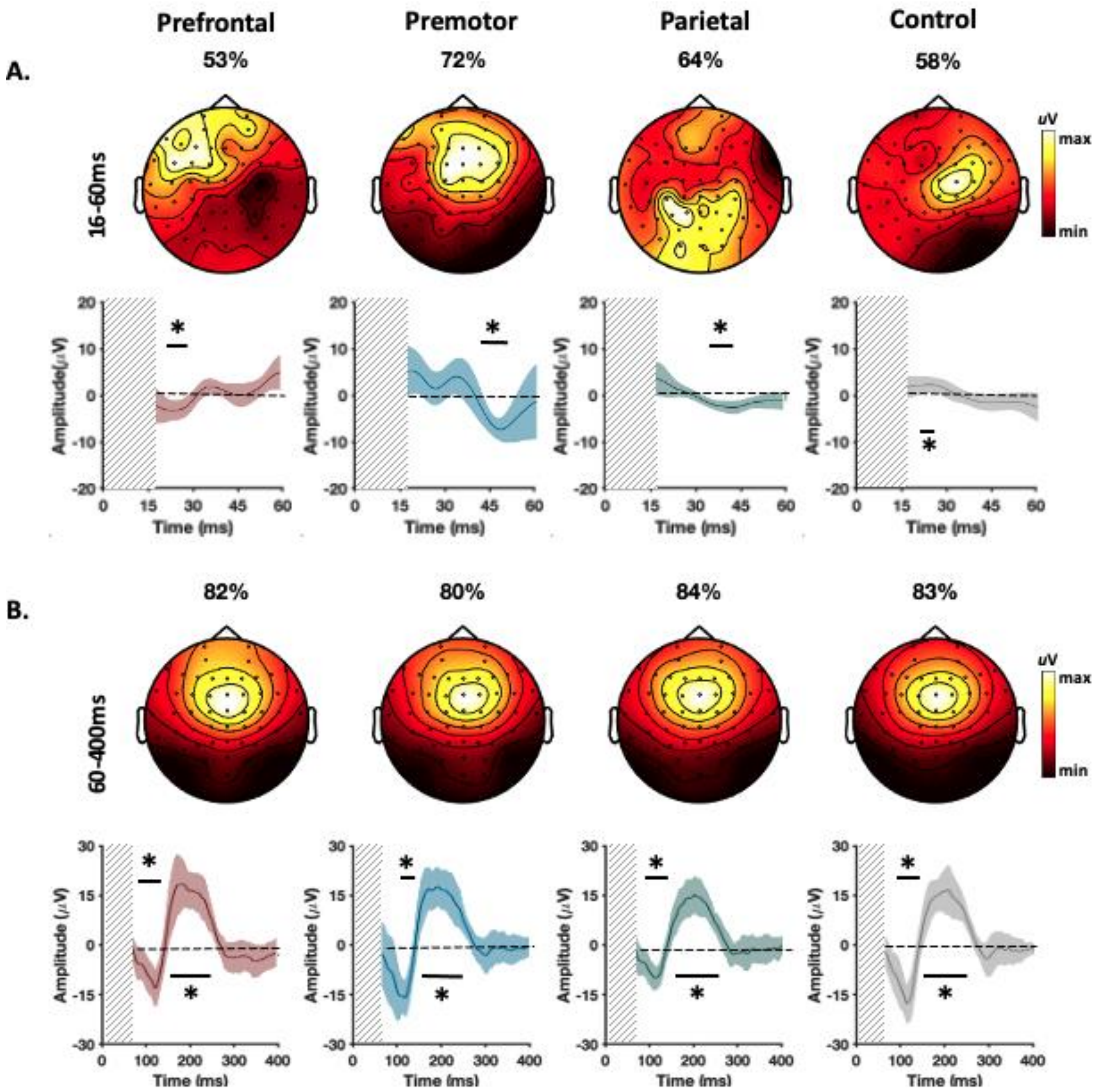


**Figure S7. The principal components that explain the maximum variance of the potentials recorded at four different stimulation conditions in experiment C.** The control condition represents TEPs from shoulder stimulation. A) The dominant PCs identified in early TEPs recorded between 16 and 60ms. B) The dominant PCs identified in late TEPs recorded between 60 and 400ms. The values above the scalp maps indicate the percentage of variance explained by the depicted component for each condition. The line graphs illustrate the changes of PCs amplitude over time. The thick lines represent the group averaged signal and the shaded areas show 95% CIs of the individual values. The vertical grey bars demonstrate the time-window of the potentials not considered for the analysis. The horizontal lines with * indicate when the signals significantly deviate from baseline (corrected $p<0.05$).
